# IMPPAT: A curated database of <u>I</u>ndian <u>M</u>edicinal <u>P</u>lants, <u>P</u>hytochemistry <u>A</u>nd <u>T</u>herapeutics

**DOI:** 10.1101/206995

**Authors:** Karthikeyan Mohanraj, Bagavathy Shanmugam Karthikeyan, R.P. Vivek-Ananth, R.P. Bharath Chand, S.R. Aparna, P. Mangalapandi, Areejit Samal

**Author notes:** M.K., B.S.K. and R.P.V. contributed equally to this work.

## Abstract

Phytochemical constituents of medicinal plants encompass a diverse space of chemical scaffolds which can be used for rational design of novel drugs. India is rich with a flora of indigenous medicinal plants that have been used for centuries in traditional Indian medicine to treat human maladies. A comprehensive online database on the phytochemistry of Indian medicinal plants will enable the application of systems biology and cheminformatic approaches towards natural product based drug discovery. In this direction, we here present, IMPPAT, a manually curated database of <u>I</u>ndian <u>M</u>edicinal <u>P</u>lants, <u>P</u>hytochemistry, <u>A</u>nd <u>T</u>herapeutics. IMPPAT contains 1742 Indian medicinal plants, 9596 phytochemicals and 1124 therapeutic uses which span across 27074 plant-phytochemical associations and 11514 plant-therapeutic associations. Notably, the curation effort led to a non-redundant *in silico* chemical library of 9596 phytochemicals with standard chemical identifiers and structure information. Using cheminformatic approaches, we have computed the physicochemical properties and drug-likeliness of the phytochemicals in IMPPAT which led to a filtered subset of 960 potential druggable phytochemicals. Moreover, a comparative analysis against FDA approved drugs suggests that majority of the druggable phytochemicals in IMPPAT are good candidates for novel prospective drugs as they have little or no structural similarity with existing drugs. The IMPPAT database is openly accessible at: https://www.imsc.res.in/~asamal/resources/imppat/home.

## Introduction

Natural products continue to play a significant role in pharmaceutical industry^1–4^ as new sources of drugs. However, recently there has been a decline in the number of marketable drugs derived from natural sources^3,4^. Furthermore, the majority of these drugs fall into already known structural scaffolds as due importance has not been given to unexplored sources of natural products for drug discovery^4^. As a result, lately, there has been significant interest in applying interdisciplinary approaches^5^ such as text mining, natural language processing (NLP)^6^, machine learning^7^, cheminformatics^8^, pharmacophore-based virtual screening^9,10^, systems biology^11,12^, systems pharmacology^13^, network pharmacology^14^ to expand the novel chemical scaffold libraries for drug discovery.

India is well known for its practice of traditional medicine and ethnopharmacology^15^. It is noteworthy that traditional Indian medicinal formulations are multi-component mixtures whose therapeutic use is based on empirical knowledge rather than a mechanistic understanding of the active ingredients in the mixture^15^. Until recently, knowledge of traditional Indian medicine including important medicinal plants and their formulations were buried within books such as Indian Materia Medica^16^ and Ayurveda Materia Medica^17^. The nondigital nature of this information limited their effective use towards new drug discovery^5^. Hence, digitization of this knowledge into a comprehensive database on Indian medicinal plants, phytochemistry and ethnopharmacology will enable researchers to apply computational approaches towards drug discovery.

Availability of a curated database of plants, their associated natural products and a repository of their chemical structures, can help in *in silico* drug discovery. In this direction, there has been significant recent progress in the development of databases^18–25^ on natural products with a focus on phytochemistry of edible and herbaceous plants. Examples of such databases include CVDHD^21^, KNAPSACK^22^, Nutrichem^18,19^, Phytochemica^20^, TCMID^23^ and TCM-Mesh^24^ which can facilitate virtual screening of prospective drug compounds or aid in the investigation of plant-disease associations. However, from the perspective of traditional Indian medicine, there have been relatively few efforts to build online databases that include Indian medicinal plants, their phytochemical constituents and therapeutic uses. Previously, Polur *et al*^26^ compiled information on 295 ayurvedic Indian medicinal plants, their 1829 phytochemical constituents and therapeutic uses. Subsequently, Polur *et al*^26^ studied the structural similarity between their library of 1829 phytochemicals and drugs in the DrugBank^27^ database to predict biologically active natural compounds. Recently, the Phytochemica^20^ database gathered information on 5 Indian medicinal plants and their 963 phytochemical constituents. In addition, Phytochemica^20^ provided chemical structures and pharmacological properties of the phytochemicals within their database. Other efforts to build online databases for traditional Indian medicine has largely been limited to cataloguing medicinal plants and their therapeutic uses rather than capturing the phytochemical constituents that are vital for drug discovery. On the other hand, in contrast to the above mentioned online databases, more comprehensive databases are available for Chinese medicinal plants. For example, TCM-MeSH^24^ is an online database for traditional Chinese medicine which captures phytochemical compositions and therapeutic uses for more than 6000 Chinese medicinal plants.

We therefore have built a manually curated database, IMPPAT, containing 1742 <u>I</u>ndian <u>M</u>edicinal <u>P</u>lants, 9596 <u>P</u>hytochemical constituents, <u>A</u>nd 1124 <u>T</u>herapeutic uses. In addition, the IMPPAT database has linked Indian medicinal plants to 974 openly accessible traditional Indian medicinal formulations. Importantly, our curation efforts have led to a non-redundant *in silico* chemical library of 9596 phytochemical constituents for which we have computed physicochemical properties using cheminformatic tools^28^. We then employed cheminformatic approaches to evaluate the drug-likeliness of the phytochemicals in our *in silico* chemical library using multiple scoring schemes such as Lipinski’s rule of five (RO5)^29^, Oral PhysChem Score (Traffic Lights)^30^, GlaxoSmithKline’s (GSK’s) 4/400^31^, Pfizer’s 3/75^32^, Veber rule^33^ and Egan rule^34^. We found a subset of 960 phytochemical constituents of Indian medicinal plants that are potentially druggable in our chemical library based on multiple scoring schemes. In summary, the IMPPAT database is a culmination of our efforts to digitize the wealth of information contained within traditional Indian medicine and provides an integrated platform where principles from systems biology and cheminformatics can be applied to accelerate natural product based drug discovery. IMPPAT is openly accessible at: https://www.imsc.res.in/~asamal/resources/imppat/home.

## Methods

### Data collection, curation and processing

**Curated list of Indian medicinal plants.** In the preliminary phase of the database construction (Figure 1), we compiled a comprehensive list of more than 5000 Indian medicinal plants based on information provided by the Foundation for Revitalisation of Local Health Traditions (FRLHT), Bengaluru (http://envis.frlht.org/), Central Institute of Medicinal and Aromatic Plants (CIMAP), Lucknow (http://cimap.res.in/) and Ministry of AYUSH, Government of India (http://ayush.gov.in/). Due to the usage of multiple synonyms for medicinal plants across sources, the common names were converted into their scientific species names and the list was manually curated to remove redundancies. The Plant List database^35^ (http://www.theplantlist.org/) was used for identifying synonyms of Indian medicinal plants.

**Figure 1:**
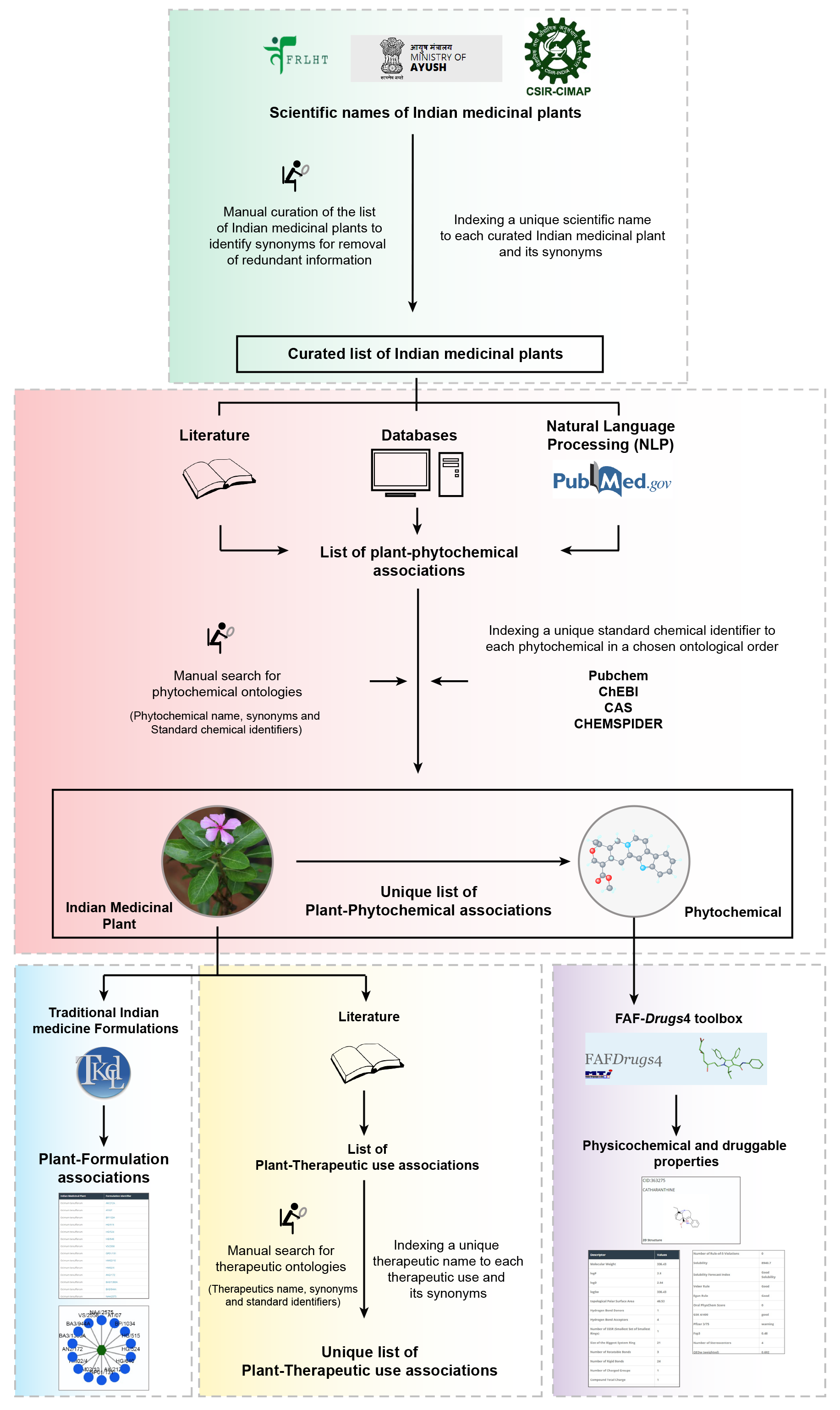
Schematic overview of the IMPPAT database construction pipeline. Briefly, we first compiled a comprehensive list of Indian medicinal plants from various sources. We next mined specialized books on Indian traditional medicine, existing databases and PubMed abstracts of journal articles to gather information on phytochemical constituents of Indian medicinal plants. We then manually annotated, curated and indexed names of identified phytochemicals with standard identifiers to build a non-redundant library of phytochemicals. This manual curation effort led to a unique list of plant-phytochemical associations. Subsequently, we gathered ethnopharmacological information from books on traditional Indian medicine to build a unique list of plant-therapeutic use associations. We also extracted publicly accessible information on traditional medicine formulations from TKDL database to build a list of plant-formulation associations. Lastly, we have used cheminformatic tools to compute the physicochemical properties and drug-likeliness of phytochemical constituents.

**Phytochemical composition of Indian medicinal plants.** After compiling a comprehensive list of more than 5000 Indian medicinal plants, we mined literature to gather information on their phytochemical constituents (Figure 1). In the first stage of data mining, we focussed on specialized traditional Indian medicine books^36–45^. From these books^36–45^, we gathered phytochemical composition for more than 1600 Indian medicinal plants. In the second stage, we gathered information from published databases of Indian medicinal plants. Phytochemica^20^ is a dedicated electronic database for the phytochemical composition of Indian medicinal plants and contains information on physicochemical properties of 963 phytochemical constituents of 5 Indian medicinal plants. Another database described in Polur *et al*^26^ had compiled information on 1829 phytochemical constituents of 295 ayurvedic Indian medicinal plants^26^. While this list is no longer publicly available, the Nutrichem^18,19^ database on phytochemical composition and therapeutic uses of plant-based food products has incorporated the information compiled by Polur *et al*^26^. From the Phytochemica^20^ and Nutrichem^18,19^ databases, we gathered information on the phytochemical composition of more than 400 Indian medicinal plants. Note that our comprehensive list covers a wide spectrum of Indian medicinal plants which includes apart from Ayurveda, other systems of traditional Indian medicine such as Siddha and Unani. In the third stage of data mining for phytochemical composition, we performed text mining of abstracts from published research articles in PubMed^46^ using natural language processing (NLP)^47^. Using in-house Python scripts, we identified keywords in PubMed abstracts which imply plant-phytochemical associations. We then used the selected keywords to mine PubMed abstracts to identify and incorporate additional references for plant-phytochemical associations in our database. In total, our database captures the phytochemical composition of 1742 Indian medicinal plants (Supplementary Table S1). The literature references for plant-phytochemical associations are listed in our database in the form of ISBN or DOI identifiers for books and PubMed identifiers (PMIDs) for journal articles.

**Annotation, curation and filtering of identified phytochemicals.** An overarching goal of this work is to create a platform for exploring the chemistry of the phytochemical constituents of Indian medicinal plants. Evaluation of the phytochemical constituents of Indian medicinal plants for their druggability or drug-likeliness will facilitate the identification of molecules for drug discovery. We would like to emphasize that synonymous chemical names are pervasive across the literature on traditional Indian medicine which were mined to construct this database. In order to remove redundancy, we manually annotated the common names of phytochemical constituents of Indian medicinal plants compiled from literature sources with documented synonyms and standard chemical identifiers (Figure 1) from Pubchem^48^, CHEBI^49^, CAS (https://www.cas.org/), CHEMSPIDER^50^, KNAPSACK^51^, CHEMFACES (http://www.chemfaces.com), FOODB (http://foodb.ca/), NIST Chemistry webbook^52^ and Human Metabolome database (HMDB)^53^. While assigning standard identifiers to phytochemicals in our database, we have chosen the following priority order: Pubchem^48^, CHEBI^49^, CAS, CHEMSPIDER^50^, KNAPSACK^51^, CHEMFACES, FOODB, NIST Chemistry webbook^52^ and HMDB^53^. We highlight that this extensive manual curation effort led to the mapping of more than 15000 common names of phytochemicals used across literature sources to a unique set of 9596 standard chemical identifiers. Phytochemicals which could not be mapped to standard chemical identifiers were excluded from our finalized database. Our choice to include only phytochemicals with standard identifiers and structure information was dictated by our goal to investigate the chemistry and druggability of phytochemical constituents of Indian medicinal plants. This largely manual effort to compile a non-redundant chemical library of 9596 phytochemical constituents of Indian medicinal plants with standard identifiers and structure information will serve as valuable resource for natural product based drug discovery in future. Moreover, the use of standard chemical identifiers will enable effortless integration of our IMPPAT database with other data sources.

**Therapeutic uses of Indian medicinal plants.** Another goal of our database is to compile ethnopharmacological information on Indian medicinal plants. Towards this goal, we manually compiled the medicinal (therapeutic) uses of Indian medicinal plants from books on Indian traditional medicine^36–45,52,54–70^. Apart from books, Polur *et al*^26^ had previously compiled a list of therapeutic uses for 295 ayurvedic Indian medicinal plants, and this information was extracted from the Nutrichem^18,19^ database. To ensure high quality, we manually curated information on therapeutic uses of Indian medicinal plants and consciously avoided automated text mining to retrieve additional information on plant-therapeutic associations. Furthermore, we manually annotated and standardized the compiled therapeutic uses of Indian medicinal plants from the above sources with identifiers from the Disease Ontology^71^, Online Mendelian Inheritance in Man (OMIM)^72^, Unified Medical Language System (UMLS)^73^ and Medical Subject Headings (MeSH)^74^ databases. To the best of our knowledge, this is the first large-scale attempt to link the ethnopharmacological information on Indian medicinal plants with standardized vocabulary in modern medicine.

**Traditional formulations of Indian medicinal plants.** Traditional knowledge digital library (TKDL) (http://www.tkdl.res.in) is a knowledgebase of traditional Indian medicinal formulations. According to TKDL, there are more than 250000 formulations of Ayurveda, Siddha and Unani of which 1200 representative formulations are openly accessible via their database. To exhibit the broader utility of our database to phytopharmacology, we have also compiled and curated the subset of 1200 openly accessible formulations in TKDL which contain at least one of the 1742 Indian medicinal plants in our database. This process led to associations between 321 Indian medicinal plants in our database and 974 traditional Indian medicinal formulations which are openly accessible through TKDL database (Figure 1). We emphasize that our database has only incorporated open digital information on traditional Indian medicinal formulations from TKDL database. However, we are aware of the vast literature^16,17,75^ on traditional Indian medicinal formulations, especially in books, and in the future, a significant effort will be needed to digitize and integrate such information into our database.

### Database management and network visualization

To construct this database, the compiled and curated data was integrated using MySQL (https://www.mysql.com/), a relational database management system which serves as a back-end for our resource. The web interface for the database was built using Drupal (https://www.drupal.org/), a PHP-based content management system hosted on Apache server with the MySQL database in the back-end. Users can browse or query our database using the scientific names of Indian medicinal plants, standard identifiers for phytochemicals, or associated therapeutic uses (Figure 2). Further we have integrated the Cytoscape.js application^76^ (http://js.cytoscape.org/) into our web interface which enables visualization of plant-phytochemical associations and plant-therapeutic associations in the form of a network. The Cytoscape network visualization displays different types of nodes such as plant, phytochemical, therapeutic use and traditional medicinal formulations in different shapes and colours. Finally, the association network can be downloaded as a tab-separated list using the available export option in our database (Figure 2).

**Figure 2:**
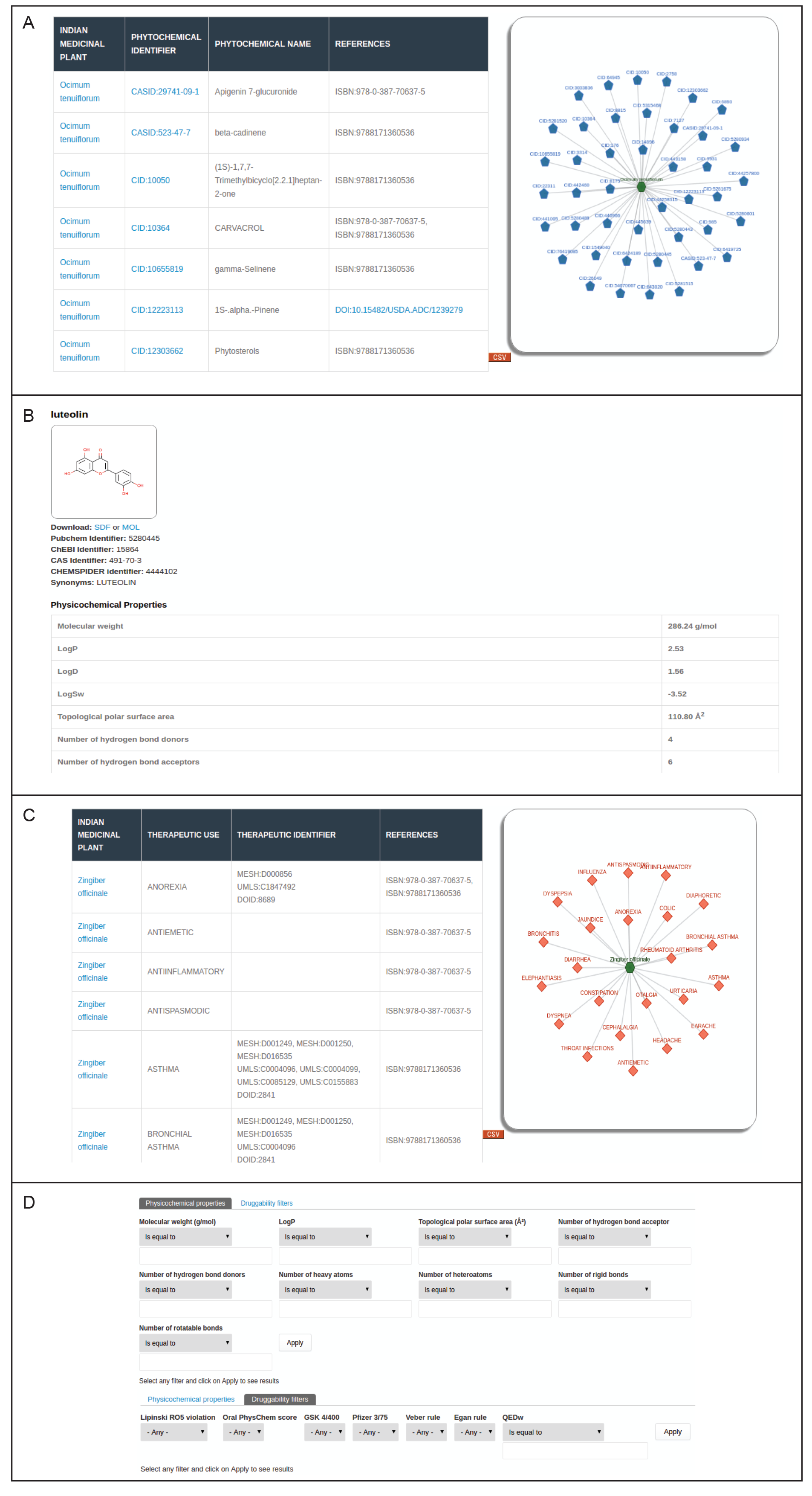
Web-interface of the IMPPAT database. (A) Snapshot of the result of a standard query for phytochemical constituents of an Indian medicinal plant. In this example, we show the plant-phytochemical association for *Ocimum tenuiflorum*, commonly known as Tulsi, from IMPPAT database. (B) Snapshot of the dedicated page containing detailed information on chemical structure, physicochemical properties and druggability scores for a chosen phytochemical. From the dedicated page for each phytochemical, users can download the structure of the phytochemical in the form of a SDF or MOL file. (C) Snapshot of the result of a standard query for therapeutic uses of an Indian medicinal plant. In this example, we show the therapeutic uses of *Zingiber officinale*, commonly known as Ginger, from IMPPAT database. (D) Snapshot of the advanced search options which enable users to filter phytochemicals based on their physiochemical properties or druggability scores.

### Computation of physicochemical properties, druggability and similarity of phytochemicals

**Physicochemical properties and druggability.** We used FAF-Drugs4 web-service^28^ to compute the following physicochemical properties of the phytochemicals: molecular weight, partition coefficient, solubility in water, topological polar surface area, charge of the compound, number of hydrogen bond donors and acceptors, number of rotatable and rigid bonds, number of hetero- and heavy atoms, and number of stereocenters. FAF-Drugs4 web-service^28^ tested the druggability of the phytochemicals based on multiple scoring schemes, namely, Lipinski’s rule of five (RO5)^29^, Oral PhysChem Score (Traffic Lights)^30^, GlaxoSmithKline’s (GSK’s) 4/400^31^, Pfizer’s 3/75^32^, Veber rule^33^ and Egan rule^34^. We filtered phytochemicals with no RO5 violation, net Traffic Lights value of zero and satisfying GSK’s 4/400, Pfizer’s 3/75, Veber rule and Egan rule as *druggable*. We further computed the weighted quantitative estimate of drug-likeness (QEDw)^77^ score using FAF-QED web-service^28^ for the filtered list of druggable phytochemicals.

**Similarity of phytochemicals.** Tanimoto coefficient (Tc)^78^ is a widely used measure to compute structural similarity between chemicals^79^. To evaluate the structural similarity of chemicals within our database to known drugs using Tc, we employed two molecular fingerprints: (a) Extended Circular Fingerprints (ECFP4)^80^ applying Morgan algorithm^81^ with radius value of 2 as implemented in RDKit^82^, and (b) MACCS keys based fingerprint. We employed the open source package, RDKit^82^, to compute molecular fingerprints and Tc between pairs of chemical structures. To identify structural similarity between chemicals, a stringent cut-off of Tc ≥ 0.5 was used while employing ECFP4 and a cut-off of Tc ≥ 0.85 was used while employing MACCS keys. Our selection of Tc cut-offs for ECFP4 and MACCS keys based computations was motivated by the recent work of Jasial *et al*^83^.

We obtained a list of 2069 FDA approved drugs from DrugBank^27^ and computed their structural similarity with our druggable phytochemicals using both ECFP4 and MACCS keys based molecular fingerprints. Note that ECFP4 molecular fingerprints were used to create the chemical similarity network of the druggable phytochemicals with QEDw score ≥ 0.9. Besides quantifying the structural similarity based on the Tc of phytochemicals, we have employed principal component analysis (PCA) to explore possible relationships between druggable phytochemicals with QEDw score ≥ 0.9 based on their physicochemical properties.

## Results

### Web-interface of the database

The IMPPAT database captures information on three types of associations for Indian medicinal plants: phytochemical composition, therapeutic uses, and traditional medicinal formulations (Figure 1). The web-interface of the database enables users to query for each of these associations using (a) scientific names of plants, (b) standard chemical identifiers of phytochemical constituents, (c) therapeutic uses, or (d) formulation identifiers (Figure 2). The web-interface displays the result of user queries for these associations in two ways: (a) A table of associations with references to literature sources, and (b) A network visualization of the associations which is powered by Cytoscape.js^76^ (Figure 2). In addition, users can also download the result of their queries for different associations of medicinal plants as a tab-separated list using the available export option in the web interface. In the results page of queries for plant-phytochemical associations, users can click each phytochemical name or identifier to navigate to a separate page containing detailed information such as chemical structure, alternate chemical names or identifiers, computed physicochemical properties, computed druggability scores and the option to download the chemical structure file in SDF format (Figure 2; Methods). Queries for plant-therapeutic associations leads to a page where users can also obtain the disease ontology identifiers corresponding to therapeutic uses (Figure 2; Methods). In the results page of queries for plant-formulation associations, users can click the medicinal formulation identifiers to navigate to the corresponding page in the TKDL database. Moreover, in the query page of IMPPAT database, users can use advanced search options (Figure 2) to filter phytochemicals based on physicochemical properties (e.g., molecular weight, number of hydrogen bond acceptors) or satisfying various druggability scores (e.g. RO5, Traffic Lights).

### Network of plant-phytochemical associations, plant-therapeutic use associations, and plant-traditional medicinal formulation associations

IMPPAT database contains information on the phytochemical composition and therapeutic uses of 1742 Indian medicinal plants (Supplementary Table S1). Of the 134 Indian medicinal plants in the priority list of Ministry of AYUSH, Government of India, 116 Indian medicinal plants are contained in our database (Supplementary Table S1). In addition, we identified 15 Indian medicinal plants in our database that appear in the red list of the International Union of Conservation of Nature (IUCN) as either near threatened, vulnerable, endangered or critically endangered (http://www.iucnredlist.org/). IMPPAT captures information on 27074 plant-phytochemical associations which encompasses 1742 Indian medicinal plants and their 9596 phytochemical constituents. Among the 1742 Indian medicinal plants in our database, *Catharanthus roseus* has the highest number of phytochemical associations. In Figure 3A, we show a histogram of the occurrence of phytochemicals across 1742 Indian medicinal plants in our database. From this figure, it is seen that the majority of phytochemicals are found in less than 5 Indian medicinal plants while only a handful of phytochemicals are found in more than 200 Indian medicinal plants.

**Figure 3:**
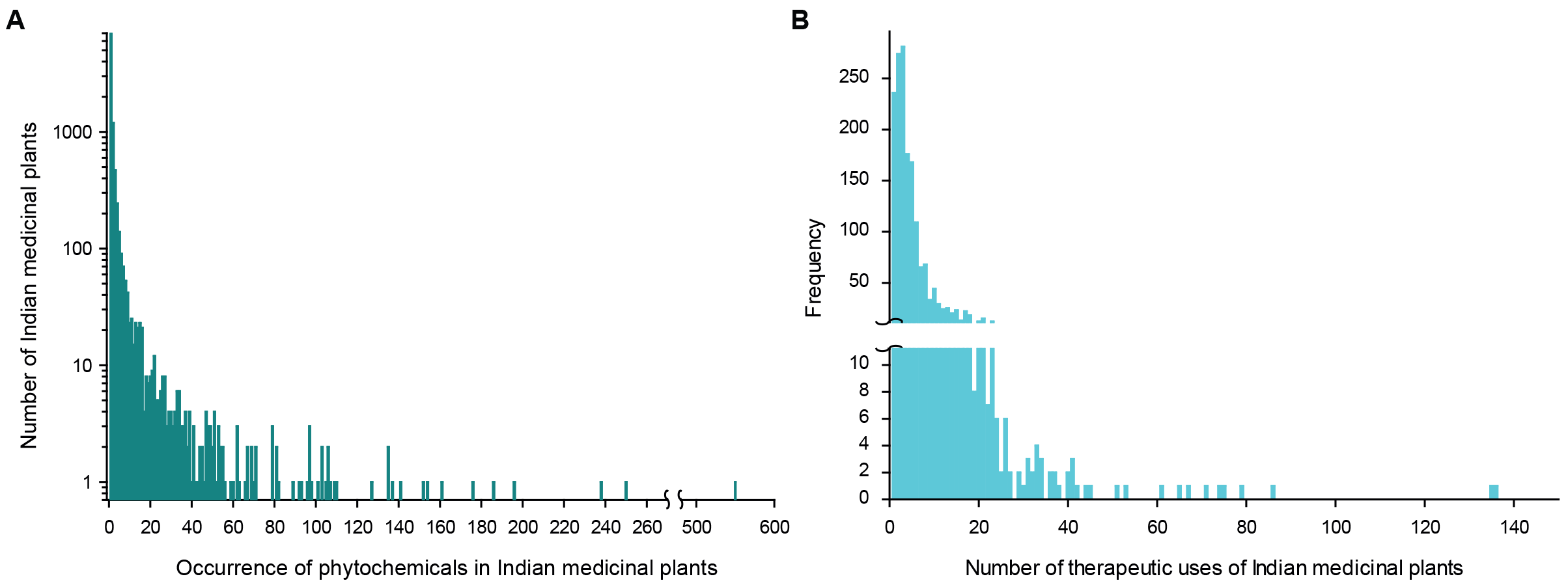
Occurrence of phytochemicals and therapeutic uses of Indian medicinal plants in IMPPAT database. (A) Histogram of the occurrence of 9596 phytochemicals across 1742 Indian medicinal plants in our database. (B) Histogram of the number of therapeutic uses per Indian medicinal plant in our database.

IMPPAT also captures information on 11514 plant-therapeutic use associations which encompasses 1742 Indian medicinal plants and 1124 therapeutic uses. In Figure 3B, we show a histogram of the number of therapeutic uses per Indian medicinal plant in our database. From this figure, it is seen that the majority of Indian medicinal plants have less than 10 documented therapeutic uses while a small fraction of Indian medicinal plants have more than 20 therapeutic uses in our database. Among the 1742 Indian medicinal plants in our database, *Ginkgo biloba*, *Panax ginseng* and *Allium sativum* have the largest number of documented therapeutic uses. Lastly, IMPPAT also captures information on 5069 plant-formulation associations which encompasses 321 Indian medicinal plants in our database and 974 traditional Indian medicinal formulations which are openly accessible from the TKDL database (Methods).

### Druggability analysis of phytochemical constituents of Indian medicinal plants

**Druggable phytochemicals.** We evaluated the druggability of 9596 phytochemicals in IMPPAT database based on multiple rules or scoring schemes, namely, RO5^29^, Traffic Lights^30^, GSK’s 4/400^31^, Pfizer’s 3/75^32^, Veber rule^33^ and Egan rule^34^ which were computed using FAF-Drugs4 web-service^28^ (Methods). The horizontal bar plot in Figure 4A gives the number of phytochemicals in IMPPAT that satisfy different druggability scores. From this figure, it is seen that the majority of our phytochemicals satisfy Veber or Egan rules in comparison to Pfizer’s 3/75 rule or net Traffic Lights value of zero. Furthermore, we find that the same set of 8712 phytochemicals in IMPPAT satisfy both the Veber rule and Egan rule for drug-likeliness. The vertical bar plot of Figure 4A shows the overlap between sets of phytochemicals that satisfy different druggability scores. We found that 960 out of 9596 phytochemicals in IMPPAT database satisfy all evaluated druggability scores (Figure 4A). Subsequently, we designated this filtered list of 960 phytochemicals as *druggable*. Among the 1742 Indian medicinal plants in our database, *Brassica oleracea*, *Catharanthus roseus*, *Zea mays*, *Oryza sativa*, *Vigna radiate*, *Pisum sativum*, *Anethum sowa*, *Allium cepa*, *Cassia obtusifolia* and *Camellia sinensis* produce the highest number of druggable phytochemicals (Table 1). In Figure 4B, we show the classification of the 960 druggable phytochemicals into broad classes similar to the classification of natural products in NPACT^25^ database. It is seen that the subset of 960 druggable phytochemicals is enriched in flavonoids and terpenoids. In Figure 4C, we show the distribution of weighted quantitative estimation of drug-likeness (QEDw) score^77^ for the 960 druggable phytochemicals. From this figure, it is seen that 14 druggable phytochemical have a QEDw score ≥ 0.9 and 98 druggable phytochemicals have a QEDw score ≥ 0.8.

**Figure 4:**
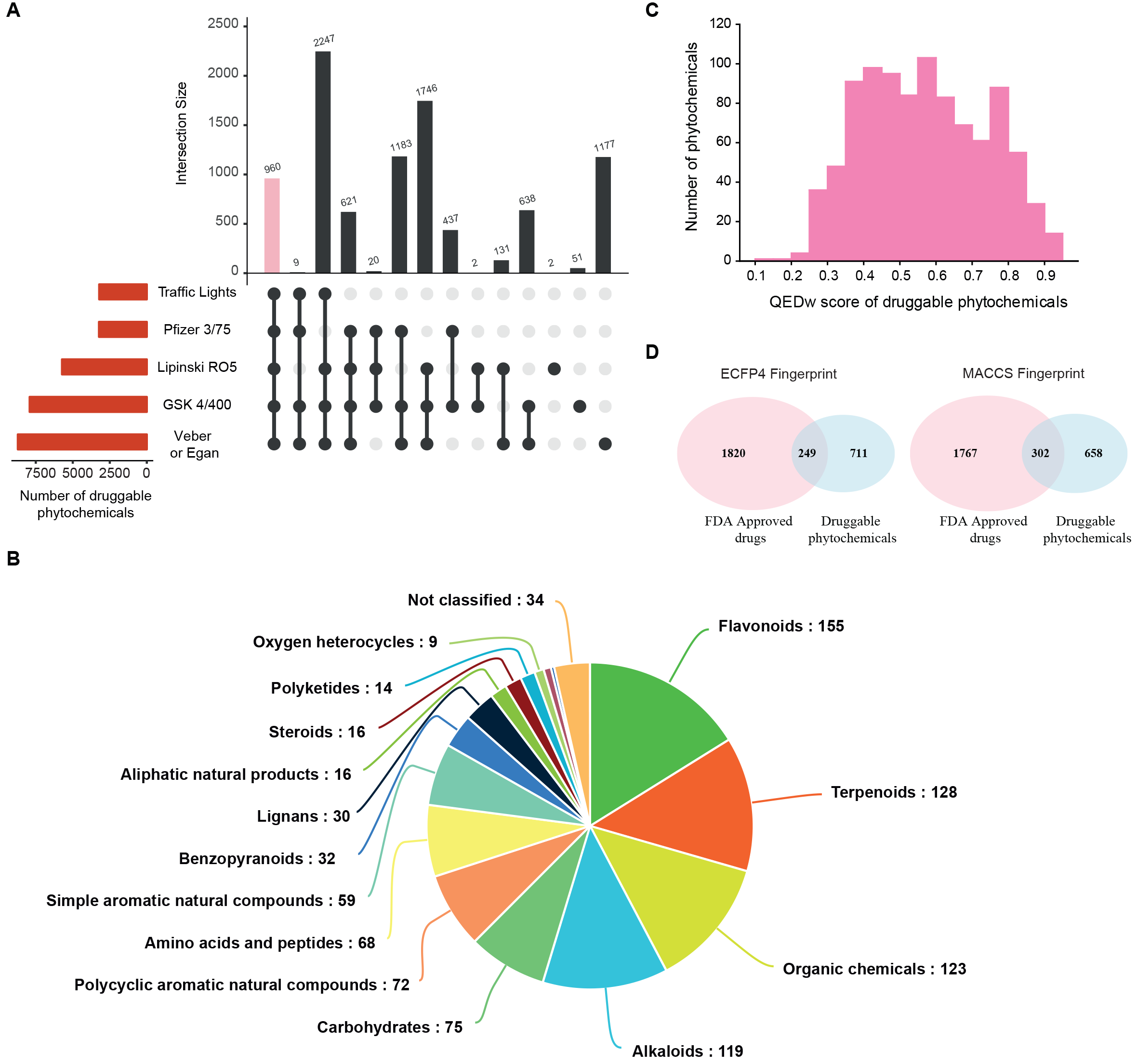
Druggability analysis of phytochemicals in IMPPAT database. (A) Evaluation of drug-likeliness of phytochemicals based on multiple scores. The horizontal bar plot shows the number of phytochemicals in the IMPPAT database that satisfy different druggability scores (Methods). The vertical bar plot shows the overlap between sets of phytochemicals that satisfy different druggability scores. The pink bar in the vertical plot gives the 960 phytochemicals which satisfy all druggability scores. This plot was generated using UpSetR^91^ package. (B) Classification of the 960 druggable phytochemicals into broad chemical classes. (C) Distribution of weighted quantitative estimate of drug-likeness (QEDw)^77^ score for the 960 phytochemicals which satisfy all druggability scores. (D) Venn diagrams summarizing structural similarity analysis of 960 druggable phytochemicals in IMPPAT database and FDA approved drugs. Based on ECFP4 and MACCS keys molecular fingerprints, 249 and 302 druggable phytochemicals, respectively, were found to be similar to FDA approved drugs.

**Table 1:**
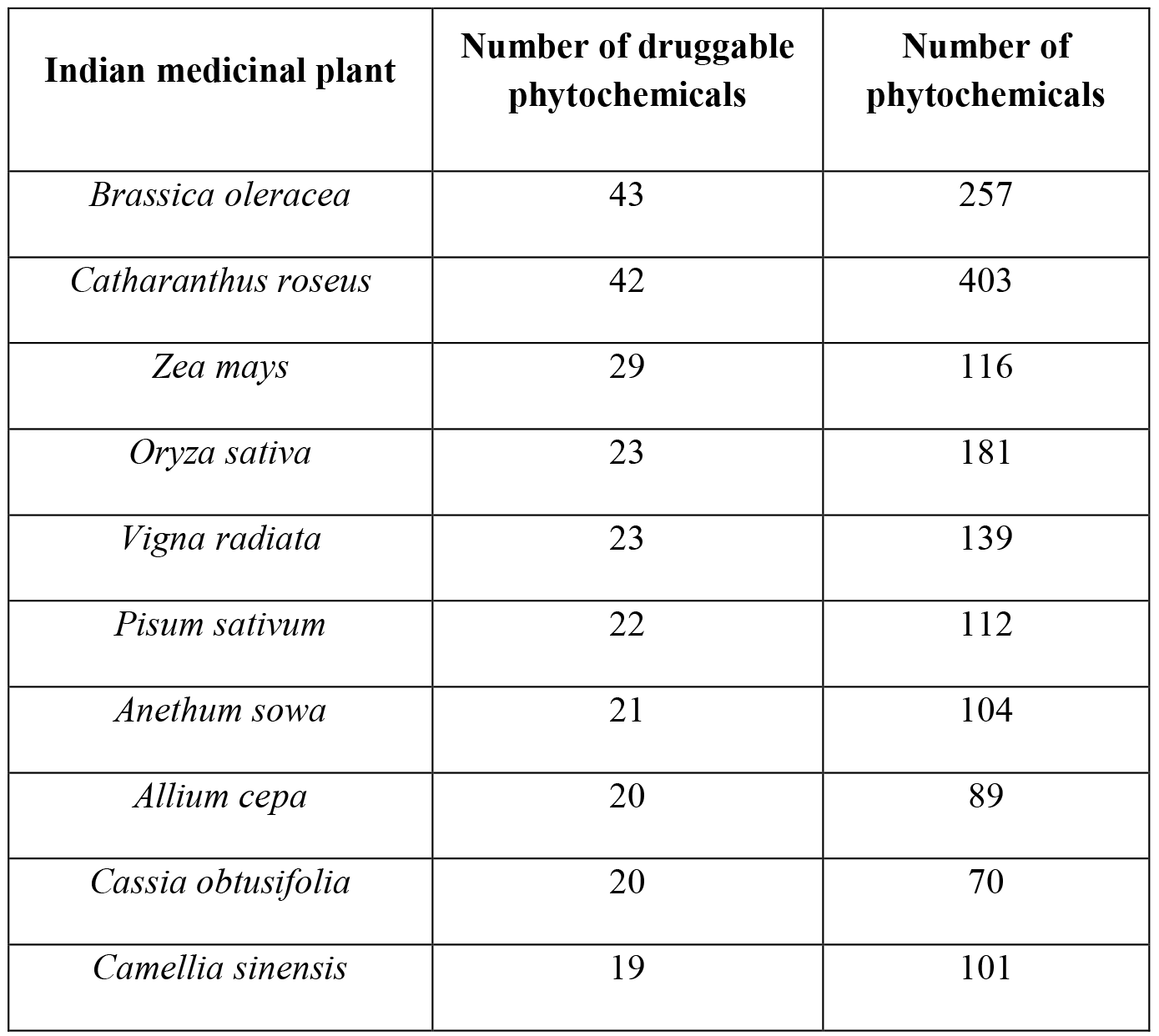
List of 10 Indian medicinal plants with the highest number of druggable phytochemicals in the IMPPAT database. The table also lists the number of phytochemical constituents of the Indian medicinal plants.

**Overlap with approved drugs space.** We obtained the structures of 2069 FDA approved drugs from the DrugBank^27^ database. By investigating the structural similarity between FDA approved drugs and 960 druggable phytochemicals in our IMPPAT database, we found that 249 and 302 druggable phytochemicals are similar to FDA approved drugs based on ECFP4 or MACCS keys molecular fingerprints respectively (Figure 4D; Methods). Combined, ECFP4 and MACCS keys based fingerprints identified 369 out of 960 druggable phytochemicals that are similar to FDA approved drugs (Methods). Thus, almost 40% of the druggable phytochemicals in IMPPAT database are similar to at least one FDA approved drug which testifies to our systemic approach to identify potential druggable phytochemical constituents of Indian medicinal plants. Importantly, the remaining 591 druggable phytochemicals which have no similarity with any of the FDA approved drugs are novel candidates for designing new drugs based on natural products from Indian medicinal plants.

**Chemical similarity network of the most-druggable phytochemicals.** For subsequent analysis, we selected 14 druggable phytochemicals with QEDw score^77^ ≥ 0.9 which were designated as the *most-druggable* phytochemicals. Of these 14 phytochemicals, 12 were found to be similar to at least one of the FDA approved drugs based on either ECFP4 or MACCS keys based molecular fingerprint. The remaining 2 most-druggable phytochemicals, Onosmone (CID:102212116) and Truxillic acid (CID:78213), were found to have no similarity with any of the FDA approved drugs. In order to probe the structural diversity of these 14 most-druggable phytochemicals, we computed the Tc based on ECFP4 molecular fingerprint between all pairs of phytochemicals (Methods). In Figure 5A, we display the similarity matrix based on Tc for the 14 most-druggable phytochemicals. From this figure, it is seen that the majority of the Tc values are small in the similarity matrix implying high structural diversity. Moreover, the similarity matrix can be transformed into a similarity network of phytochemicals by using a stringent threshold value of Tc ≥ 0.5 to determine edges in the graph (Figure 5B). We find that only 16 of the 91 possible edges between the 14 most-druggable phytochemicals are realized in the similarity network (Figure 5B). Furthermore, the similarity network can be partitioned into a large connected component (cluster) of 7 phytochemicals, a smaller connected component of 2 phytochemicals and 5 remaining isolated phytochemicals. We highlight that the 2 phytochemicals, Onosmone and Truxillic acid, that have no similarity with any of the FDA approved drugs are among the isolated nodes in the similarity network (Figure 5B). Based on plant-phytochemical associations in our database, Onosmone and Truxillic acid are phytochemical constituents of Indian medicinal plants, *Onosma echioides* and *Erythroxylum coca*, respectively, and a survey of the literature shows that these phytochemicals are under active investigation for their therapeutic uses^84–88^. We also highlight that none of the 14 most-druggable phytochemicals are captured by Phytochemica^20^ database while 6 of the 14 phytochemicals are captured by Nutrichem^18,19^ database.

**Figure 5:**
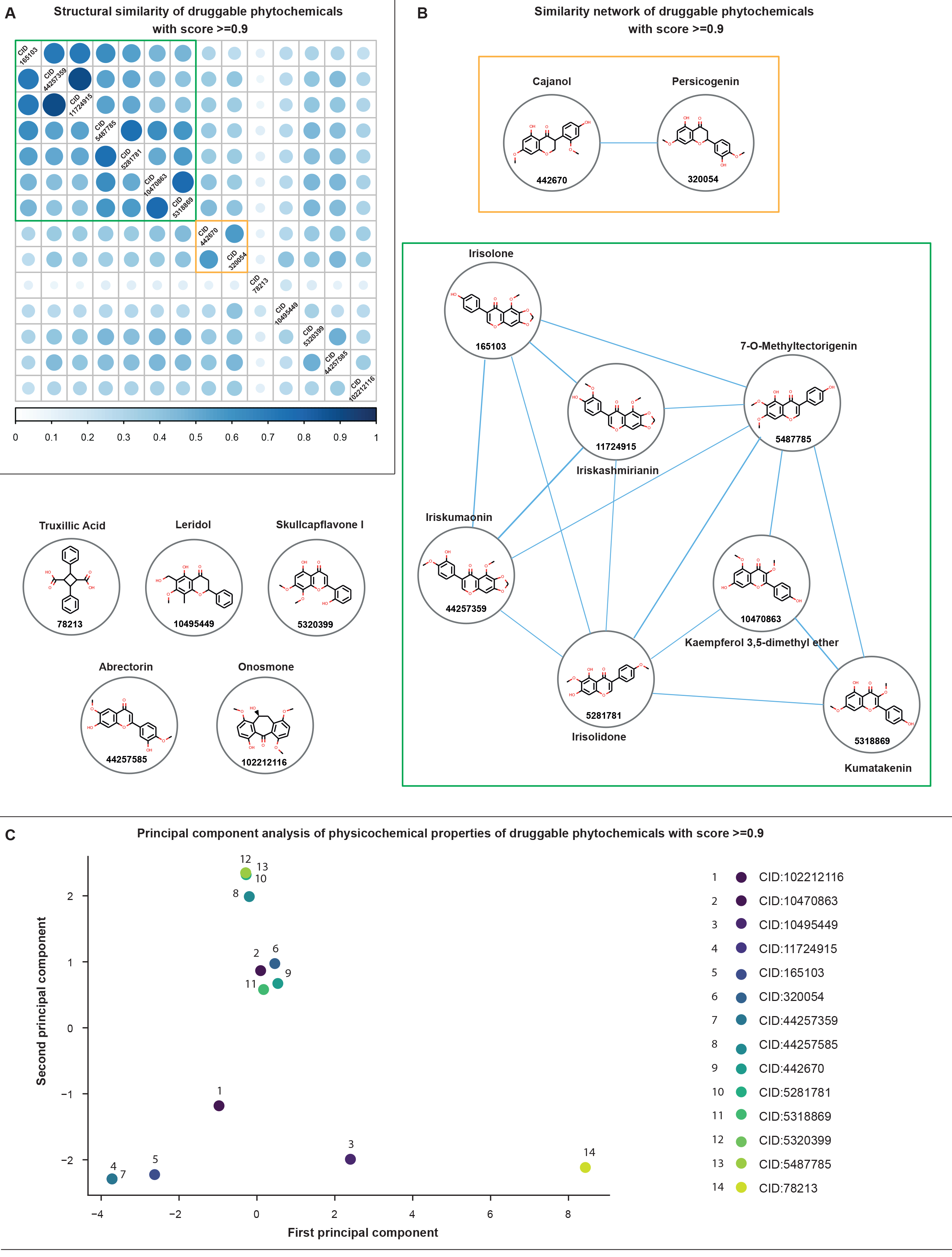
Structural similarity and physicochemical properties of most-druggable phytochemicals in IMPPAT database. (A) Similarity matrix for the 14 most-druggable phytochemicals with QEDw score ≥ 0.9 based on Tanimoto coefficient (Tc) between pairs of chemicals computed using ECFP4 molecular fingerprints. (B) Similarity network for the 14 most-druggable phytochemicals constructed using a stringent threshold value of Tc ≥ 0.5 to determine edges in the graph. We find that the similarity network can be partitioned into a large connected component of 7 phytochemicals, a smaller connected component of 2 phytochemicals and 5 isolated phytochemicals. (C) Principal component analysis (PCA) of the 14 most-druggable phytochemicals based on their physicochemical properties. The first and second principal components can together explain more than 71% of the total variance in the dataset.

**Principal component analysis of the most-druggable phytochemicals based on their physicochemical properties.** In the last section, we studied the similarity between chemical structures of 14 most-druggable phytochemicals to find a large cluster of 7 phytochemicals with highly similar chemical structures. But it is well known that high similarity between chemical structures does not necessarily imply high similarity between chemical activities^89^. Thus, we here investigate the physicochemical properties of the 14 most-druggable phytochemicals. Note that we have used FAF-Drugs4 web-service^28^ to compute several physicochemical properties including molecular weight, partition coefficient, solubility in water, topological polar surface area, charge of the compound, number of hydrogen bond donors and acceptors, number of rotatable and rigid bonds, number of hetero- and heavy atoms, and number of stereocenters for each phytochemical in IMPPAT database (Methods). In Figure 6, we show the distribution of four physicochemical properties, namely, molecular weight, number of hydrogen bond donors, number of hydrogen bond acceptors and topological polar surface area, across the 9596 phytochemicals in our database. We performed principal component analysis (PCA) of the 14 most-druggable phytochemicals based on their physiochemical properties (Figure 5C). In Figure 5C, the first and second principal components together explained more than 71% of the total variance in the dataset. We find that the 7 most-druggable phytochemicals which are clustered together in the structural similarity space (Figure 5B) are not clustered together in the physicochemical or chemical activity space (Figure 5C). These observations based on limited analysis of 14 most-druggable phytochemicals suggest that a combined exploration of chemical similarity space and physicochemical or chemical activity space of phytochemical constituents of Indian medicinal plants will be required in the future to identify and design novel drugs.

**Figure 6:**
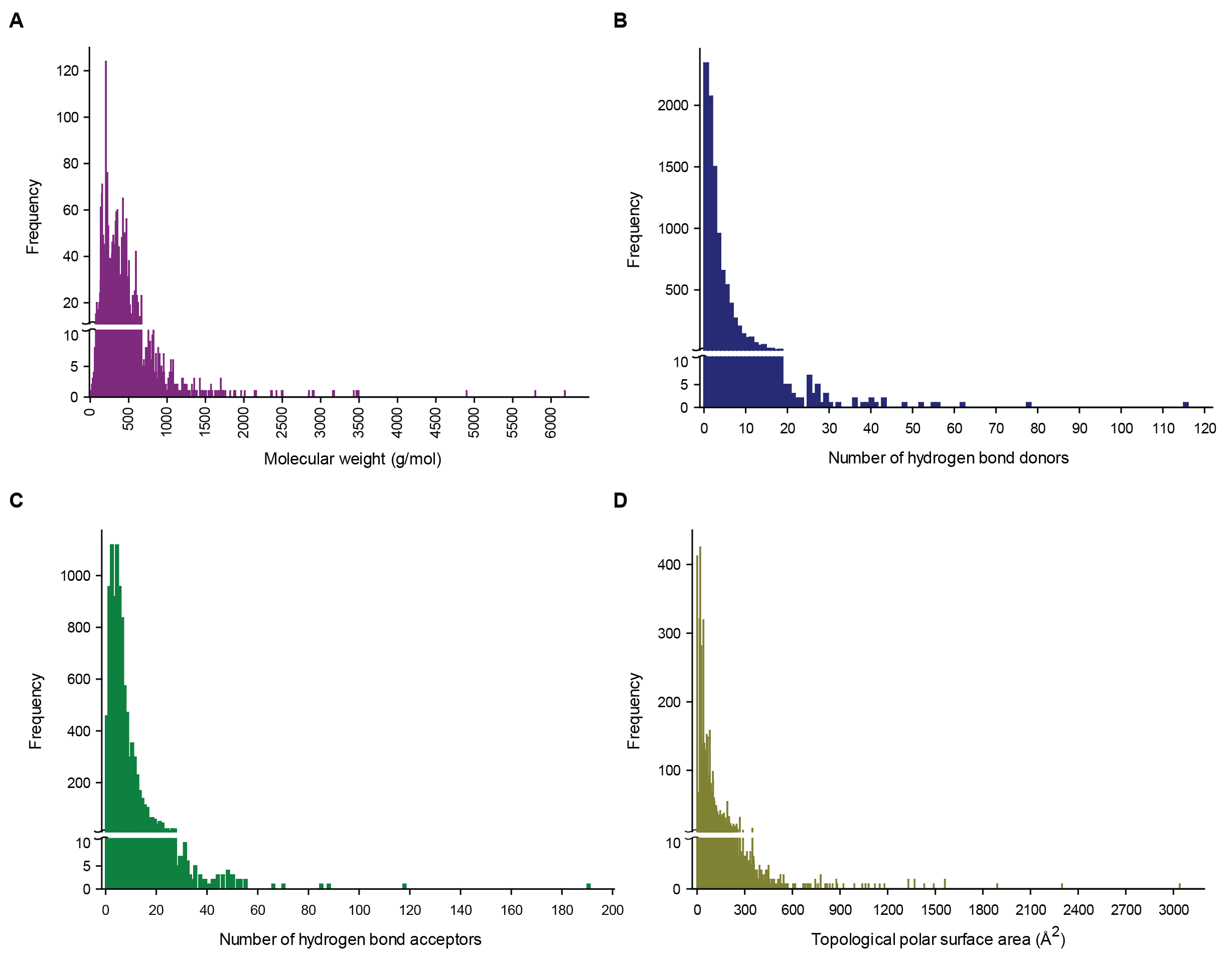
Distribution of physicochemical properties across the 9596 phytochemicals in IMPPAT database. (A) Molecular weight. (B) Number of hydrogen bond donors. (C) Number of hydrogen bond acceptors. (D) Topological polar surface area.

### Discussion and future directions

In the 21^st^ century, there is immense interest within academia and pharmaceutical industry to incorporate systems biology approaches to accelerate the drug discovery pipeline which has led to the emergence of sub-disciplines such as systems pharmacology^13^ and network pharmacology^14^. Likewise cheminformatics can accelerate drug discovery by aiding in the rational design of robust chemical scaffolds from diverse natural sources^5^. Recently, Lagunin *et al*^5^ have reviewed more than 50 existing bioinformatics resources for natural product based drug discovery. In their review, Lagunin *et al*^5^ noted that most existing databases on medicinal plants and associated phytochemistry do not provide a platform to integrate systems biology and cheminformatics approaches which impedes natural products based drug discovery. Towards this goal, we here incorporate principles from systems biology and cheminformatics to build an extensive resource on phytochemistry and ethnopharmacology of Indian medicinal plants. Here we present, IMPPAT, a curated database of 1742 <u>I</u>ndian <u>M</u>edicinal <u>P</u>lants, 9596 <u>P</u>hytochemical constituents, <u>A</u>nd 1124 <u>T</u>herapeutic uses. Importantly, IMPPAT provides a unifying platform for the application of system-level approaches to elucidate mechanistic links between phytochemical constituents of Indian medicinal plants and their therapeutic action.

Using cheminformatic approaches, we found that 960 of the 9596 phytochemical constituents of Indian medicinal plants in our database are potentially druggable based on multiple scoring schemes. Of the 960 phytochemicals which satisfy all druggability scores evaluated here, a subset of 14 phytochemicals were found to have a QEDw score^77^ ≥ 0.9 (Figure 5). Interestingly, the occurrence of these 14 most-druggable phytochemicals across 1742 Indian medicinal plants in our database is very rare with none of the 14 phytochemicals being found in more than 3 Indian medicinal plants. Specifically, the 14 most-druggable phytochemicals are constituents of only 17 Indian medicinal plants in our database. Also, 4 of the 14 most-druggable phytochemicals are constituents of 3 phylogenetically close Indian medicinal plants, *Iris germanica, Iris nepalensis* and *Iris kemaonensis*, which are from the same genus. However, we find that only 2 out of 17 Indian medicinal plants that produce the 14 most-druggable phytochemicals are in the priority list of Ministry of AYUSH, Government of India. This analysis suggests a possible revision in the AYUSH priority list to include the remaining 15 Indian medicinal plants that produced the majority of the most-druggable phytochemicals in our database. Thus, our resource will facilitate rational design of scaffolds for new drugs based on natural products and future expansion of the chemotaxonomy^90^ of Indian medicinal plants.

In the future, we hope to update IMPPAT database with the following additional information. Firstly, it will be important to link the phytochemical constituents of the Indian medicinal plants with their gene or protein targets. Such target information is vital to obtain a mechanistic understanding of either the therapeutic action or toxic effects of Indian medicinal plants. For example, TCM-Mesh^24^, a traditional Chinese medicine database has gathered gene or protein target information for phytochemical constituents of Chinese medicinal plants using a network pharmacology approach. Importantly, information on gene or protein targets of phytochemicals will also enable pathway level assessment of the therapeutic action of medicinal plants which will help design robust drug scaffolds for many complex diseases. Secondly, it will be important to update our database with more detailed information on the parts of the Indian medicinal plants such as leaves, stem or root, that produce the different phytochemical constituents. Such detailed information on the phytochemical composition of parts of Indian medicinal plants will be crucial for evaluating and developing traditional Indian medicine formulations^75^. However, significant manual curation and literature mining will be needed to expand our database to include the phytochemical composition of the different parts of 1742 Indian medicinal plants which is beyond the scope of the present work. Thirdly, it will be important to enrich our database by incorporating more traditional Indian medicinal formulations. For example, TKDL (http://www.tkdl.res.in) has made only 1200 of their documented 250000 traditional Indian medicinal formulations openly accessible, and future efforts to associate this wealth of information to our database will shed mechanistic information on the therapeutic action of traditional formulations. Lastly, it will be important to perform a comparative analysis of the phytochemical composition of Indian medicinal plants with those of Chinese medicinal plants. Such a comparative analysis will shed light on phytochemicals and druggable scaffolds exclusive to Indian medicinal plants.

## Acknowledgments

We would like to thank Gopal C. Nanda and the staff of Achanta Lakshmipathi Research Centre for Ayurveda, Chennai for discussions and help in accessing relevant literature. We also thank S. Krishnaswamy and James Craig for discussions and the library staff of The Institute of Mathematical Sciences, Chennai for facilitating access to scientific literature. AS acknowledges financial support from Department of Science and Technology (DST) India through the award of a start-up grant (YSS/2015/000060) and Ramanujan fellowship (SB/S2/RJN-006/2014), Max Planck Society Germany through the award of a Max Planck Partner Group, and Department of Atomic Energy (DAE) India. The funders have no role in study design, data collection, data analysis, manuscript preparation or decision to publish.

## Author contributions

A.S., B.S.K., R.P.V. and M.K. designed research. M.K., B.S.K., R.P.V, R.P.B., S.R.A. and P.M. compiled and curated data from various sources. M.K. designed the database platform and visual interface. R.P.V. performed the cheminformatic analysis. B.S.K., R.P.V. and A.S. wrote the manuscript. All authors have read and approved the manuscript.

## Supplementary Material

**Supplementary Table S1:** List of 1742 Indian medicinal plants with information on the phytochemical composition and therapeutic uses in IMPPAT database. Of the 1742 Indian medicinal plants in IMPPAT database, 116 are on the priority list of Ministry of AYUSH, Government of India and 15 are on the red list of the International Union of Conservation of Nature (IUCN).

